# Anti-COVID-19 Activity of FDA Approved Drugs through RNA G-quadruplex Binding

**DOI:** 10.1101/2022.05.31.493843

**Authors:** Shuvra Shekhar Roy, Shalu Sharma, Zaigham Abbas Rizvi, Dipanjali Sinha, Divya Gupta, Mercy Rophina, Paras Sehgal, Srikanth Sadhu, Manas Ranjan Tripathy, Sweety Samal, Souvik Maiti, Vinod Scaria, Sridhar Sivasubbu, Amit Awasthi, Krishnan H Harshan, Sanjeev Jain, Shantanu Chowdhury

## Abstract

The COVID-19 pandemic caused by SARS-CoV-2 has caused millions of infections and deaths worldwide. Limited treatment options and the threat from emerging variants underline the need for novel and widely accessible therapeutics. G-quadruplexes (G4s) are nucleic acid secondary structures known to affect many cellular processes including viral replication and transcription. We identified heretofore not reported G4s with remarkably low mutation frequency across >5 million SARS-CoV-2 genomes. The G4 structure was targeted using FDA-approved drugs that can bind G4s - Chlorpromazine (CPZ) and Prochlorperazine (PCZ). We found significant inhibition in lung pathology and lung viral load of SARS-CoV-2 challenged hamsters when treated with CPZ, PCZ that was comparable to the widely used antiviral drug Remdesivir. In support, in vitro G4 binding, inhibition of reverse transcription from RNA isolated from COVID-infected humans, and attenuated viral replication and infectivity in Vero cell cultures were clear in case of both CPZ/PCZ. Apart from the wide accessibility of CPZ/PCZ, targeting relatively invariant nucleic acid structures poses an attractive strategy against fast mutating viruses like SARS-CoV-2.

## Introduction

Since the emergence of SARS-CoV-2 more than 500 million infections and more than 6 million deaths have been confirmed (https://covid19.who.int/, as of 31^st^ May, 2022). Recommended treatments for COVID-19 include immunomodulatory molecules like JAK inhibitors, IL-6 receptor blockers, systemic corticosteroids; monoclonal antibodies targeting the spike protein of SARS-CoV-2; and drugs like Remdesivir, Nirmatrelvir/ritonavir and Molnupiravir that target viral replication (WHO; Janik et al., 2021; Kelleni, 2021). However, these are not readily accessible in a global context underlining a critical need for developing both effective and affordable therapeutic strategies against SARS-CoV-2.

Sequences within viral genomes adopt secondary structures that can be targeted using ligands as potential antiviral drugs ^3–6^. DNA or RNA sequence motifs with four or more runs of guanine repeats interspersed with short runs of other bases form non-canonical secondary structures called G-quadruplexes (G4s) (reveiwed in Mukherjee et al., 2019; Sengupta et al., 2020; Varshney et al., 2020). Further, G4s were shown to be predominant in gene regulatory regions across organisms through specific G4-binding transcription factors ^10–14^; and, RNA G4s within mRNA were shown to inhibit translation by stalling or dissociation of ribosomes ^15–18^. Notably, recent studies identified potential G4s (pG4) in the SARS-Cov2 genome ^5,19–21^.

To target SARS-CoV-2 we sought to test FDA-approved drugs reported to bind to G4s ^22,23^. Here we show two such drugs chlorpromazine (CPZ) and prochlorperazine (PCZ) ^22^ decrease infectivity, viral load, and pathology of SARS-CoV-2 infection in the hamster model. This was further supported with intracellular results using the Vero cell infection model. Both CPZ and PCZ bound to SARS-CoV-2 RNA G4 and inhibited reverse transcription suggesting attenuated viral replication and transcription within the Vero cells in presence of CPZ/PCZ. Together these support the function of CPZ and PCZ as molecules with anti-COVID-19 activity with potential for repurposing as affordable drugs against SARS-CoV-2.

## Results

### G4 motifs in the SARS-CoV-2 genome remarkably resistant to mutations

The SARS-CoV-2 genome sequence was examined for potential G4-forming (pG4) sequence. We identified two pG4s having three G-quartets within the ORF1a/ORF1ab gene (pG4-1 and pG4-2; Table 1) not reported earlier. The pG4s included breaks within the G-stretches thus allowing bulge formations for stable G4 structure formation as deduced from multiple published G4-stability scoring algorithms (Table S1) ^24–28^. In an earlier genome-wide study in humans we noted pG4s to be relatively impervious to mutations compared to other regions of the genome, supporting physiological relevance of pG4s ^29^. Here we reasoned, if functionally significant the newly identified pG4s might be relatively less prone to mutations. Importantly, if so, such pG4s could be crucial for any approach that targets different variants of SARS-CoV-2.

**Table 1:**
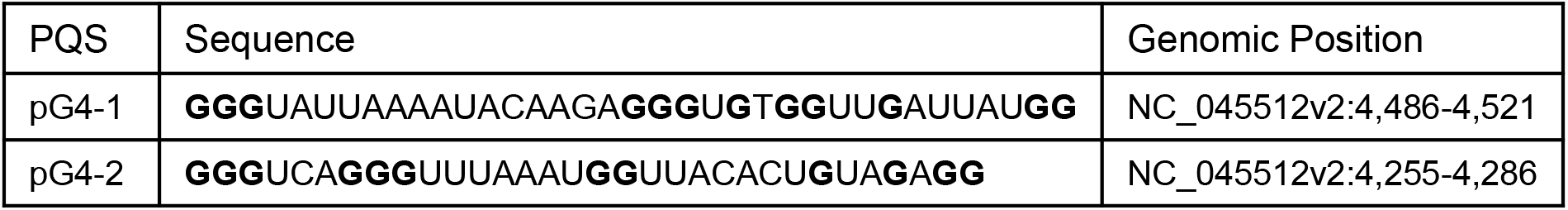
New pG4s identified in the SARS-CoV-2 genome that have bulges. Their sequences, genomic positions are mentioned.

Reported mutations within more than 5 million SARS-CoV-2 genomes (5406687 genome sequences retrieved from China National Center for Bioinformation-National Genomics Data Center as of 28^th^ of May, 2022) was analysed: On average each nucleotide was mutated ∼158 times per 100000 sequenced genomes (frequency: 0.00158; Figure 1a). The average mutation frequency of the Gs within the newly identified pG4s was significantly lower: ∼22 and 44 per 100000 SARS-CoV-2 genomes (frequency: 0.00022 and 0.00044) for pG4-1 and pG4-2, respectively (Figure 1b, c).

**Figure 1:**
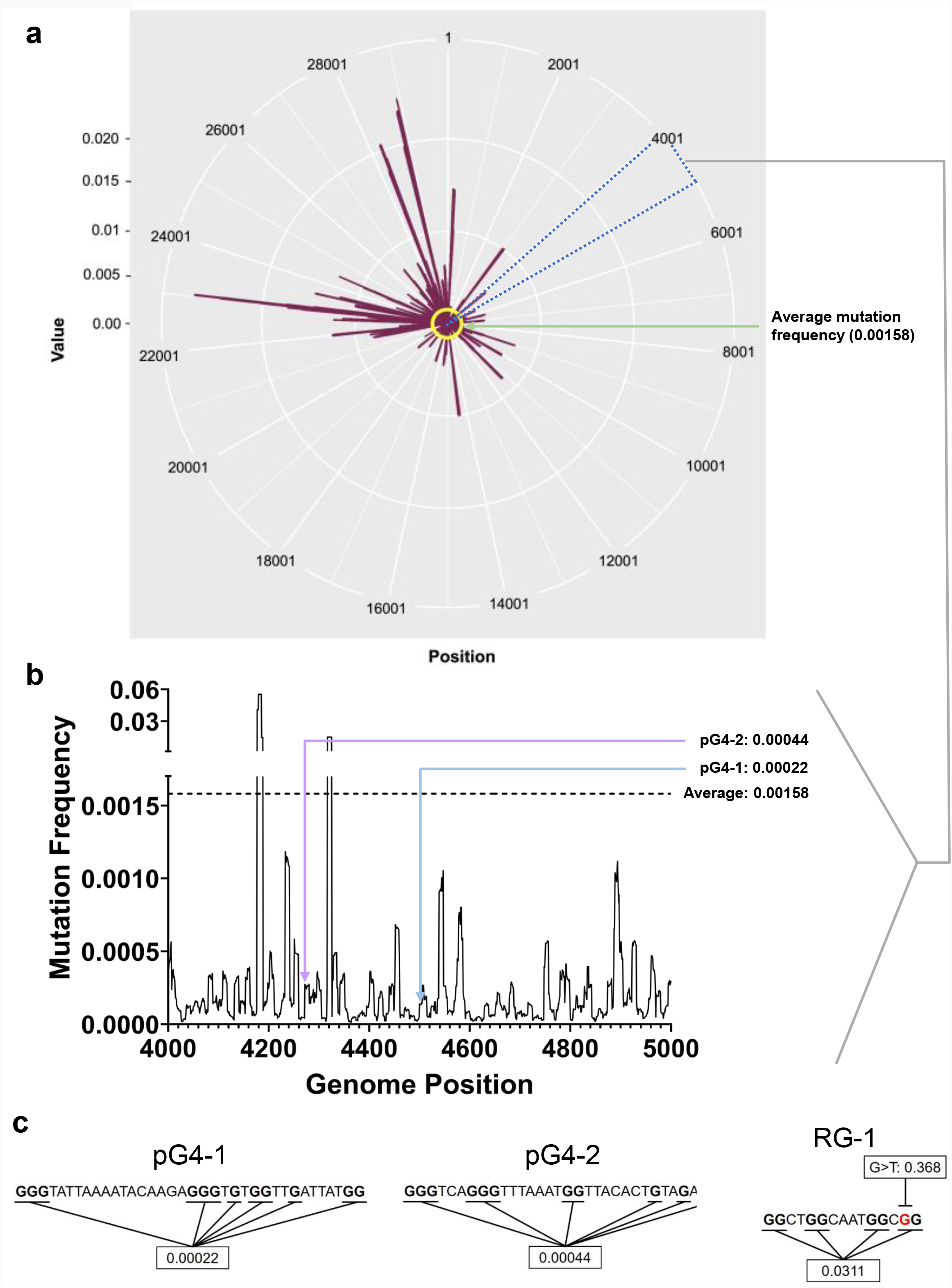
**a** Mutation frequency (number of sequenced genomes with mutation at a given base position/total number of sequenced genomes) of each nucleotide of the SARS-CoV-2 genome plotted against their respective genomic positions. The average mutation frequency is marked in yellow. **b** Mutation frequency within the 4000-5000 bp positions to clearly denote the mutation of pG4-1 and pG4-2. **c** The sequences of pG4-1, pG4-2 and RG-1 with the Gs used to calculate the average mutation frequency marked in bold. In RG-1, one of the Gs has a very high mutation frequency for a G to T mutation (marked in red).

We also checked the mutation frequency of another previously well characterised pG4 with two G-tetrads designated as RG-1 by authors ^30^. The Gs in RG-1 pG4 had a relatively high mutation frequency of 0.0311 as a constituent G was highly mutated across samples (Figure 1a, c). Based on these we further focused on pG4-1, which appeared to be most conserved, for further experiments.

### ORF1 pG4-1 forms the G4 structure

Circular Dichroism (CD) spectrum of the RNA sequence representing pG4-1 showed positive/negative peaks at 260/240 nm respectively confirming the formation of a G4 structure with parallel topology (Figure 2a) ^31^. In contrast, for a similar RNA sequence where specific Gs were substituted (mutated-control RNA), the positive/negative CD peaks were significantly attenuated supporting specificity of the folded RNA G4 (Figure 2a).

**Figure 2:**
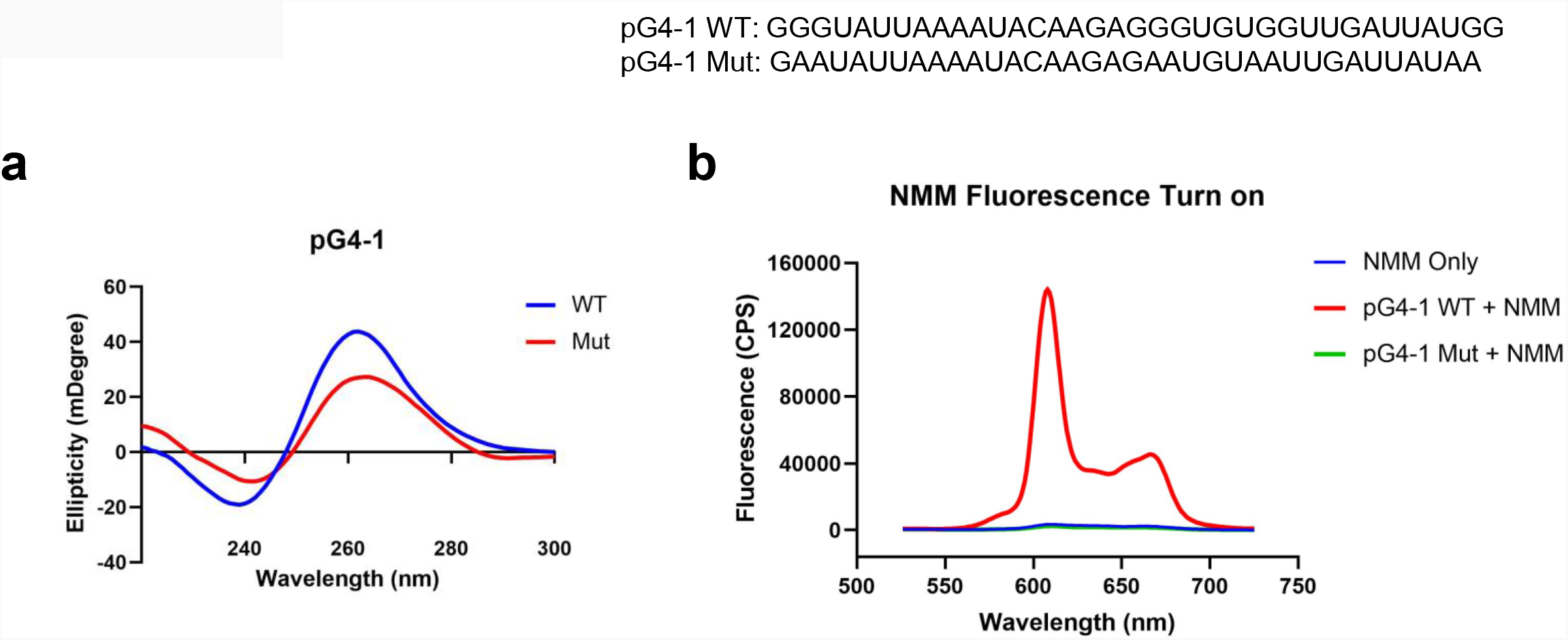
**a** CD spectra of 5 μM of pG4-1 WT and its mutated control, pG4-1 Mut where specific ‘G’s were mutated to ‘A’s to prevent G4 formation. **b** Fluorescence spectra of 1 μM NMM only or in presence of 1 μM pG4-1 WT or pG4-1 Mut.

Formation and stability of the G4 was also tested using N-methyl mesoporphyrin IX (NMM), a G4 binding ligand that gives enhanced fluorescence upon binding specifically to G4s ^32^. The fluorescence of NMM was enhanced by more than hundred-fold in presence of the pG4-1, but not in case of the mutated-control RNA sequence (Figure 2b). Together these support RNA G4 formation by the pG4-1 RNA sequence.

### CPZ and PCZ bind to ORF1 pG4-1

To design a small-molecule induced targeting we next asked if pG4-1 binds to the G4 binding ligands CPZ and/or PCZ, which are FDA approved drugs ^22,33^. Isothermal titration calorimetry (ITC) showed both CPZ and PCZ interact with pG4-1 with dissociation constants in the micromolar range: The binding between CPZ and pG4-1 was monophasic (K_d_ = 39.0 ± 1.78 μM), whereas PCZ binds to pG4-1 in a biphasic manner (K_d1_ = 32.1 ± 5.56 μM, K_d2_ = 39.0 ± 1.78 μM) (Figure 3a, b). CD of pG4-1 was performed in presence of increasing concentrations of the ligands CPZ and PCZ to test whether interaction with the ligands affected the topology of pG4-1. Results showed reduced amplitude of the positive/negative CD peaks without any change in the position of the CD peak confirming interaction of pG4-1 with CPZ and PCZ retained the parallel topology (Figure 3c, d).

**Figure 3:**
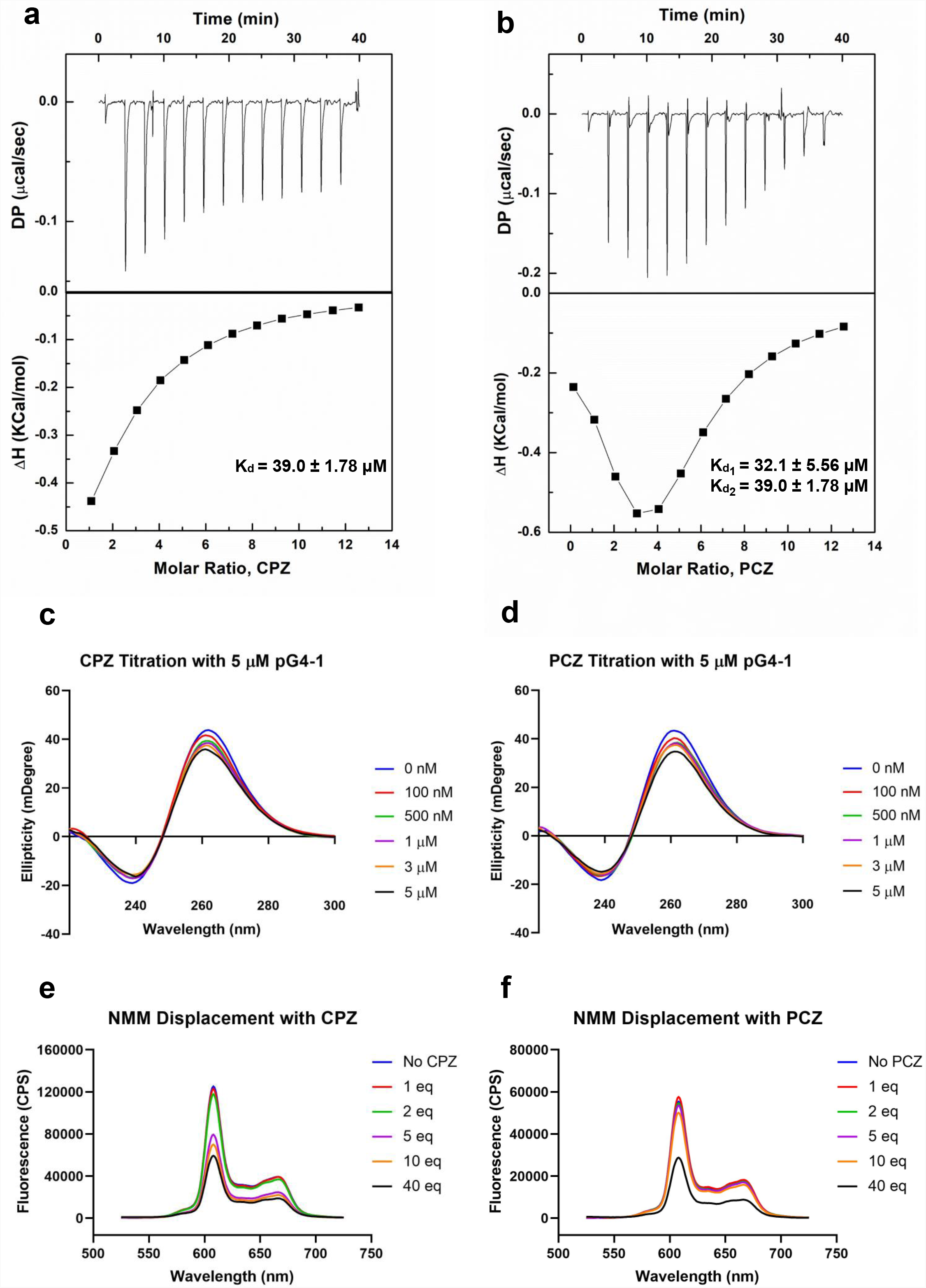
Isothermal calorimetry (ITC) profiles for the titration of CPZ (**a**) or PCZ (**b**) with ORF1 pG4-1. The upper panels show sequential injection of the ligands into ORF1 pG4-1; fitted data of integrated heat values after correction for heat of dilution plotted against molar ratio shown in lower panels; dissociation constants mentioned in the lower panel. CD spectra of ORF1 pG4-1 with increasing concentration of CPZ (**c**) or PCZ (**d**). Fluorescence spectra of 1 μM of NMM in presence of 1 μM of ORF1 pG4-1 with increasing molar ratio of CPZ (**e**) or PCZ (**f**).

We reasoned that specific binding of CPZ and PCZ with pG4-1 would compete with G4-bound-NMM resulting in reduced fluorescence from G4-NMM interaction (see above). Increasing concentration of CPZ and PCZ resulted in reduction of fluorescence from G4-NMM with more than fifty percent decrease at a molar ratio of 1:40. Together these supported specific association of both CPZ and PCZ with the RNA G4 formed by ORF1 pG4-1 (Figure 3e, f).

### CPZ and PCZ attenuate reverse transcription of SARS-CoV-2 RNA

Stabilization of RNA G4s can inhibit mammalian translation as well as replication and transcription of some viruses, via stalling and/or dissociation of polymerases or ribosomes ^15– 17,34–36^. Here we tested whether binding of CPZ/PCZ to RNA G4 affected reverse transcription. For this a synthetic SARS-CoV-2 genomic RNA template was obtained where the whole viral genome was represented by six non-overlapping 5 kb fragments. This template was reverse transcribed using specific primers such that ORF1 pG4-1 or a non-G4 forming control region was reverse transcribed. Following this, specific primers encompassing the pG4-1 or the selected control region were used for quantitative RT-PCR to assay the levels of reverse transcription (see Methods). In presence of both CPZ and PCZ reverse transcription of the pG4-1-harbouring region was significantly reduced relative to untreated samples, when normalized with the control region. Together these showed reduced reverse transcription from the pG4-1-harbouring region in presence of CPZ or PCZ (Figure 4a).

**Figure 4:**
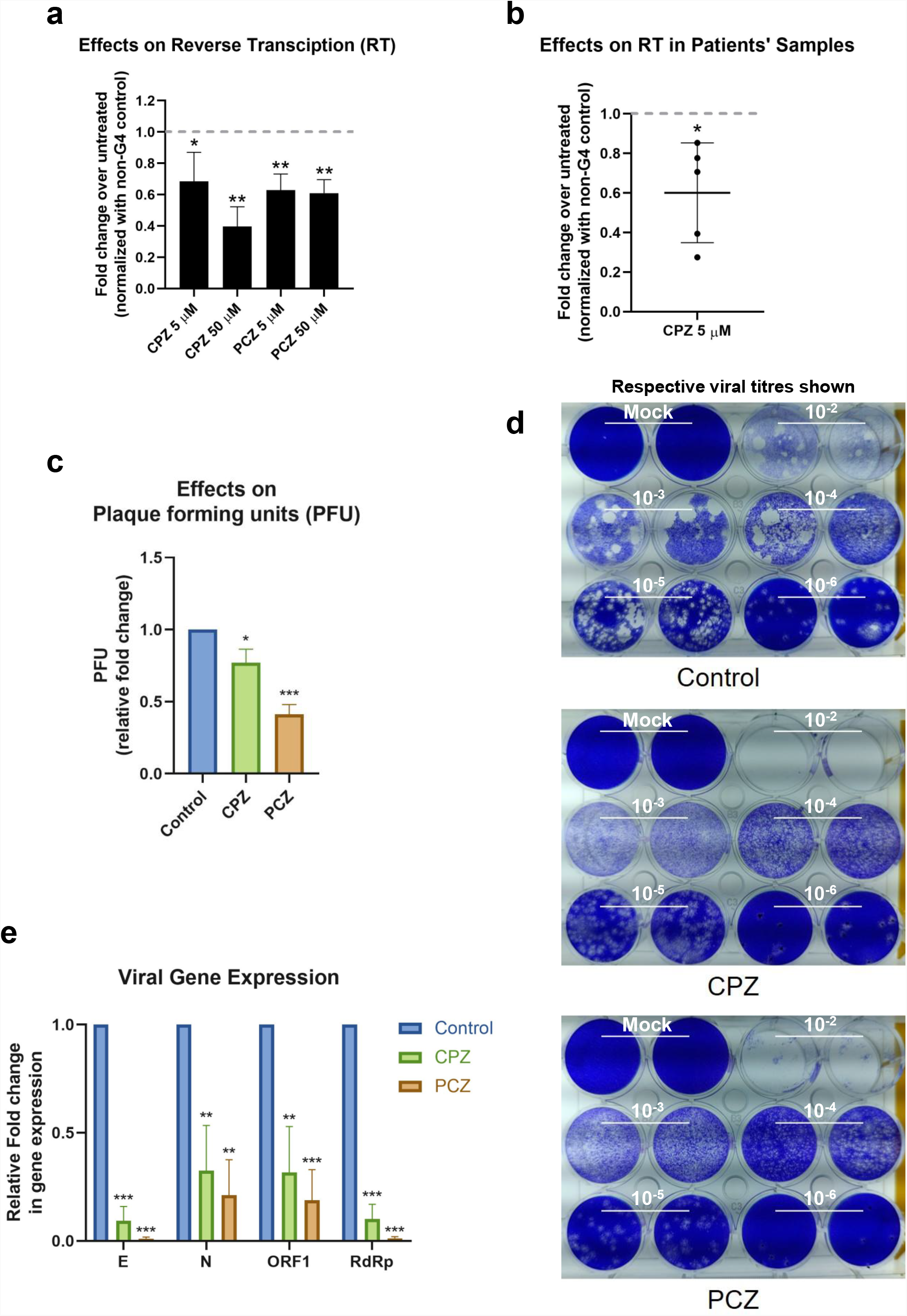
**a** Fold change in reverse transcription of the ORF1 pG4-1 harbouring region over untreated control using a synthetic SARS-CoV-2 RNA: different concentrations of CPZ and PCZ were used; a non-G4 forming region was used for normalization. Mean ± SD (n=3); unpaired, two-tailed *t*-test. **b** Fold change in reverse transcription of the ORF1 pG4-1 harbouring region over untreated control in RNA isolated from nasopharyngeal swabs of COVID-19 infected individuals, treated with 5 μM CPZ; a non-G4 forming region was used for normalization. Mean ± SD (n=3); unpaired, two-tailed *t*-test. **c** Fold change in plaque formation units (PFUs) indicating infectivity of supernatant from infected Vero cells treated with 5 μM of CPZ or PCZ compared against untreated control. Mean ± SD (n=3); unpaired, two-tailed *t*-test. **d** Representative PFU assay plates showing consistent drop in the plaques with logarithmic dilution of the sample and the effects on the number of PFUs after CPZ or PCZ treatment. Respective viral titres shown above the wells. **e** Expression of E, N, ORF1 and RdRp genes of SARS-CoV-2 in the extracellular media of infected Vero cells treated with 5 μM of CPZ or PCZ compared against untreated control. Mean ± SD (n=3); unpaired, two-tailed *t*-test.

To test in human patient samples, we used RNA isolated from nasopharyngeal swabs of five COVID-19 infected individuals. Due to the paucity of patient sample RNA only CPZ was tested. Notably, inhibition of reverse transcription in all the five samples was clear in presence of 5 micromolar CPZ (Figure 4b) suggesting interaction of CPZ with pG4-1 restricts processivity of the reverse transcriptase suggesting this might affect other translocating complexes/enzymes like RNA dependent RNA polymerase (*RdRp*).

### CPZ and PCZ can inhibit SARS-CoV-2 infectivity in Vero cells

Next, we sought to test whether CPZ and PCZ affect the infectivity of SARS-CoV-2 and viral replication within cells. Vero cells were infected with SARS-CoV-2 virus at 1 MOI for 3 h at 37°C in the presence of the respective inhibitors CPZ or PCZ either at 2µM or 5µM. After the inoculum was removed and the cells were washed, treatment was continued in 10% FBS medium for another 24 h. At 24 hpi the media was replaced by fresh media containing the respective inhibitors (Figure S1). At 48 hpi supernatants were collected for plaque formation assay and viral RNA quantification (Figure S1).

Treatment with 5 μM of CPZ or PCZ led to significant decrease in plaque formation, 25 % and 60 % for CPZ and PCZ respectively, and therefore infectivity of the virus (Figure 4c, d). The levels of viral RNA genes are indicative of viral replication and reproduction after infection. Real-time PCR quantification of the *RdRp* (RNA-dependent RNA polymerase), *E* (envelope), *N* (Nucleocapsid) and *ORF1* genes were done using RNA isolated from the collected supernatants. At 5 μM, both CPZ and PCZ treatment reduced expression of *RdRp, E, N* and *ORF1* by more than 60% (Figure 4e). However, we noted 5 μM PCZ inhibited growth of Vero cells (MTT assay; Figure S2a), whereas treatment with a lower concentration of 2 μM PCZ did not result in cytotoxicity while still reducing expression of the viral *RdRp* and *E* genes (Figure S2b, S3).

### CPZ and PCZ treatment alleviates COVID-19 pathogenesis in hamsters

To check the effect of the G4 binding ligands as potential molecules for therapeutic intervention in COVID-19 treatment we used SARS-CoV-2 infection in hamsters as previously described ^37–40^. Both prophylactic and therapeutic regimens of CPZ and PCZ treatment were tested. Prophylactic groups of hamsters received 8mg/kg or 5mg/kg of CPZ and PCZ (mentioned as pCPZ and pPCZ to denote prophylactic arm of the experiment) respectively through intraperitoneal administration each day starting from 3 days prior to the SARS-CoV-2 challenge till the end point (day 4 post infection). In the therapeutic group CPZ and PCZ (tCPZ and tPCZ to denote therapeutic arm), the hamsters received the drug 8 mg/kg and 5 mg/kg respectively through intra-peritoneal injection every day from the day of challenge till the end point ^41,42^. FDA approved drug Remdesivir, which has been shown to work as a potent antiviral against SARS-CoV-2 infection, was used as a control with 15 mg/kg subcutaneous injections given 1 day before and after the challenge ^39,43,44^.

From day 2 onwards, we noted the treatment groups had protection against body weight loss compared to the infected control group (Figure 5a). Hamsters receiving prophylactic treatment of CPZ showed no body weight loss and the trend in percent body weight change was comparable to the Remdesivir control group. Notably, prophylactic treatment of CPZ showed improved rescue of body weight loss than the prophylactic PCZ group, as well as their respective therapeutic groups, suggesting CPZ pre-treatment might be more effective in rescuing body weight loss in SARS-CoV-2 challenged hamsters (Figure 5a, b).

**Figure 5:**
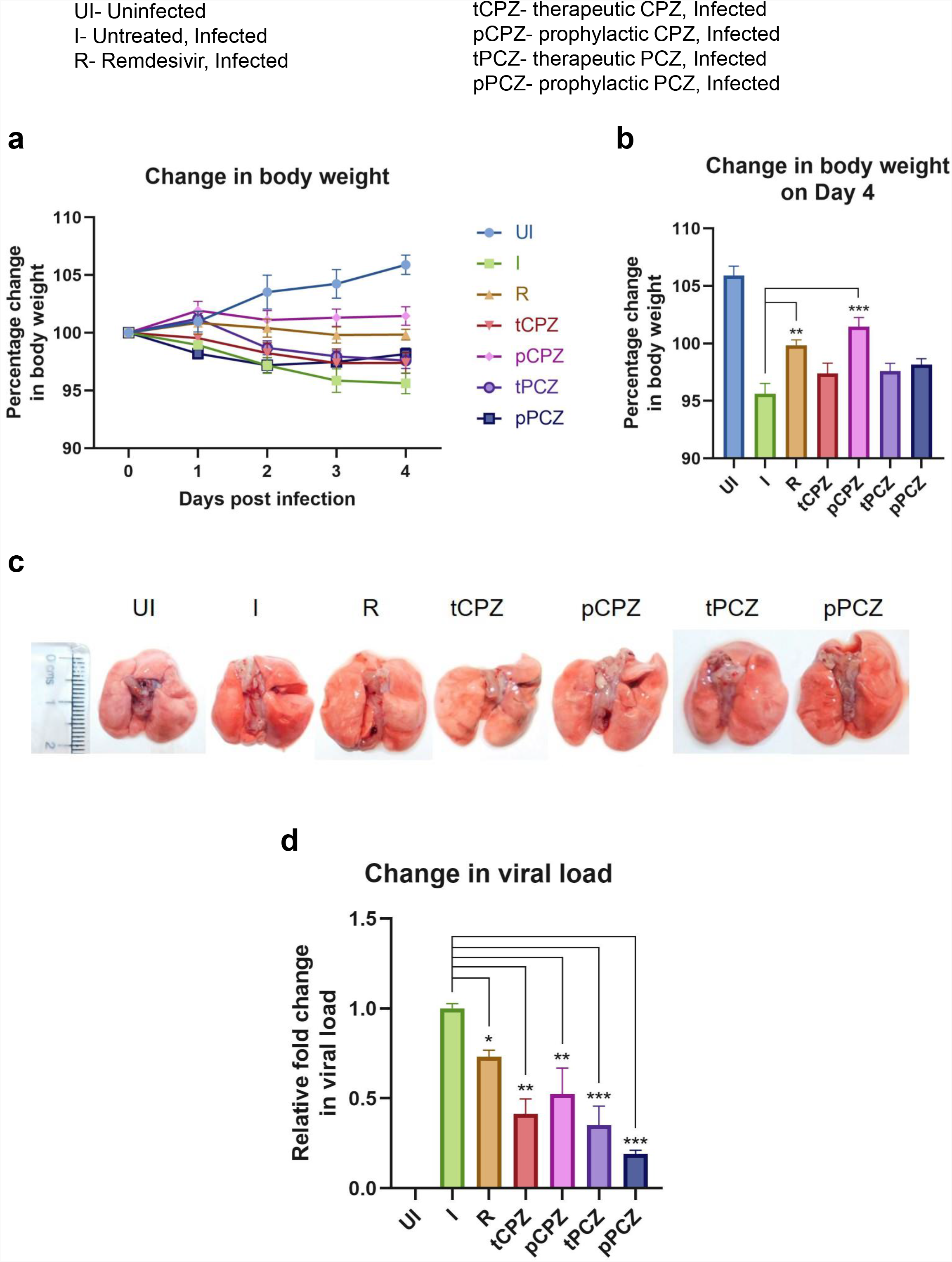
**a** Percentage (%) change in body weight of hamsters of different treatment groups: uninfected (UI), infected (I), infected with Remdesivir treatment (R), infected with prophylactic treatment of CPZ (pCPZ) and PCZ (pPCZ), infected with therapeutic treatment of CPZ (tCPZ) and PCZ (tPCZ). **b** Percentage % change in body weight of hamsters of different treatment groups on the fourth day post infection. Mean ± SE (n=3); ordinary one-way ANOVA. **c** Images of the excised lungs from hamsters of different treatment groups, showing gross morphology with pneumonitis region. **d** Lung viral load of the hamsters of different treatment groups quantified using a kit targeting the N gene of SARS-CoV-2. Mean ± SE (n=3); ordinary one-way ANOVA.

All the animals were euthanized on day 4 post challenge, which has been shown earlier as the optimal time point to study SARS-CoV-2 lung pathology in hamsters ^39,40^. Regions of pneumonia observed in the lungs of the infected animals were significantly reduced in drug-treated group in a manner similar to the remdesivir control group (Figure 5c). In order to further understand the anti-viral efficacy of the G4 binding drugs, lung viral load qPCR was performed (Figure 5d). This showed significant decrease in lung viral load in both PCZ and CPZ groups. Remarkably, when compared to the Remdesivir control group PCZ as well as CPZ administration, both as prophylactic or therapeutic drug, showed similar or more decrease in viral load as seen with the Remdesivir treatment validating the potent anti-viral nature of these drugs.

To further understand the mitigation in SARS-CoV-2-mediated lung pathology in CPZ/PCZ treated groups we carried out lung histopathological analysis through hematoxylin and eosin (H and E)–staining. The treated groups showed reduction in pneumonitis, bronchitis and inflammation similar to that of the Remdesivir control group (Figure 6a, b). In addition, we noted that the difference in alveolar epithelial injury was not significantly different in any of the drug treated groups compared to infected untreated control. Considering the overall disease score, both CPZ and PCZ could significantly reduce the lung pathology arising from the infection in a manner similar to the Remdesivir control group. Furthermore, spleen size of the animals were compared to evaluate gross morphological changes as splenomegaly can be a critical indicator of active infection ^39,40^. Importantly, the CPZ/PCZ treated groups showed reduced splenomegaly similar to Remdesivir treatment (Figure 6c).

**Figure 6:**
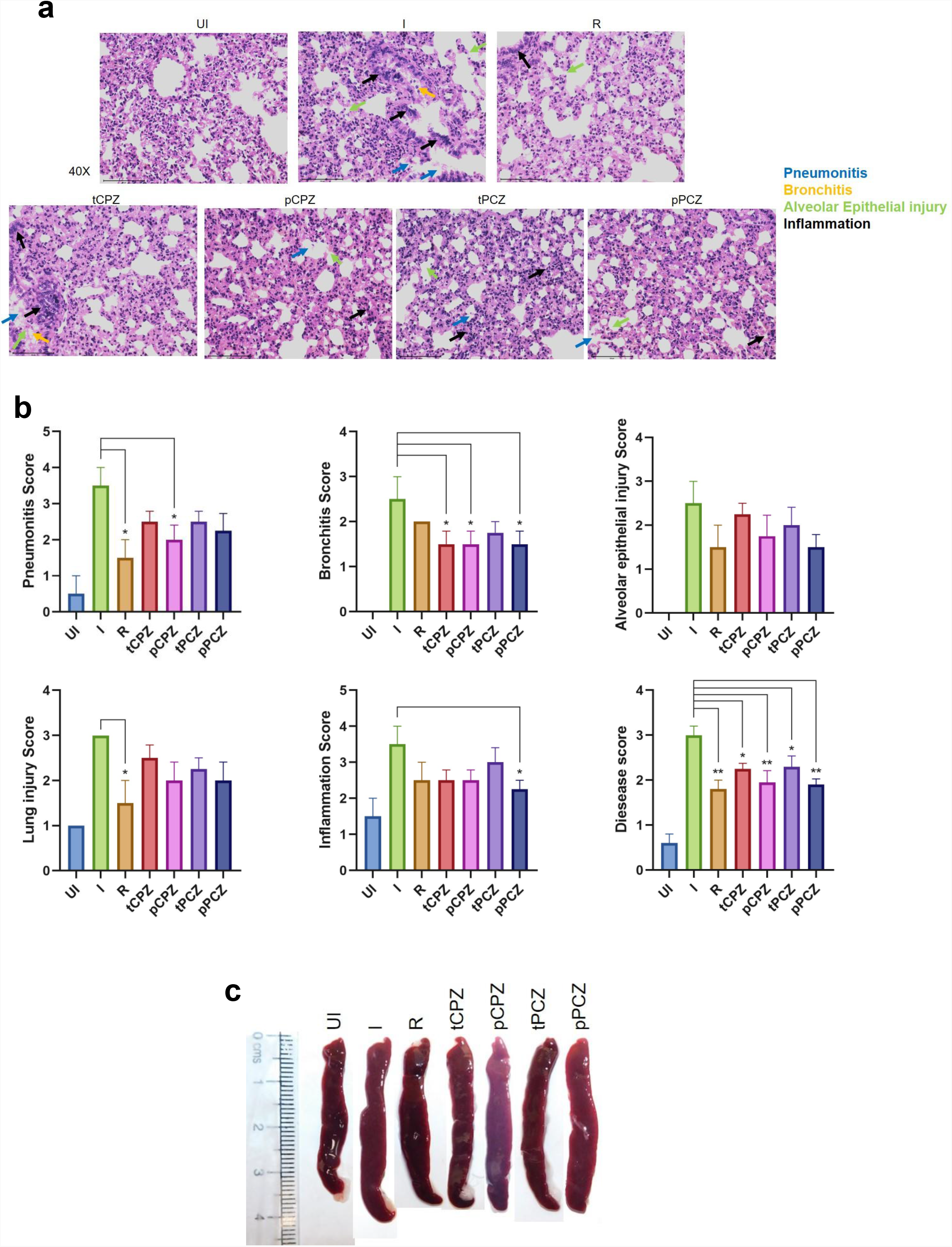
**a** Images of H&E-staining of excised lungs from hamsters of different treatment groups at ×40 magnification showing regions of pneumonitis (blue arrow), bronchitis (yellow arrow), alveolar epithelial injury (green arrow), and inflammation (black arrow). **b** Histological score for pneumonitis, bronchitis, alveolar epithelial injury, lung injury, inflammation, and overall disease score of the lungs after different treatments. Mean ± SE (n=3); ordinary one-way ANOVA. **c** Images of excised spleen indicating changes in the spleen length upon different treatments.

## Discussion

While focusing on canonical G4s to target in the SARS-CoV-2 genome we first noted that the pG4s identified so far harboured two G-tetrads (i.e., with repeats constituting 2Gs that form 2 guanine-tetrads). Typically, these tend to be less stable than G4s made of three G-tetrads ^27,45^. Here we identified two new pG4s with three G-tetrads that included bulges – a form of G4 that has been widely studied and understood to be stable in solution ^8,9^. On genome wide mutation frequency analysis using more than five million reported SARS-CoV-2 genomes we found, surprisingly, that the G4s were also significantly resistant to mutations. Selecting the more conserved of the two G4s we showed that it can form a stable G4 structure. CPZ and PCZ, two FDA approved drugs, bind to the G4 with micromolar affinity, and inhibited reverse transcription from synthetic as well as RNA isolated from patients suggesting potential anti-COVID activity. In Vero cells CPZ and PCZ inhibited SARS-CoV-2 infection and also decreased viral replication and transcription. Prompted by these results we asked whether CPZ and/or PCZ could affect infection in the hamster animal model: Notably, CPZ and PCZ decreased disease pathogenesis in terms of body weight loss, lung viral load, lung histopathology and splenomegaly. The effects of CPZ and PCZ were remarkably comparable to that of treatment with the standard-of-care antiviral Remdesivir.

Earlier studies suggested the significance of RNA G4s in infectious human viruses, and a few recent reports focused particularly on G4s in SARS-CoV-2 ^5,19–21^. Nsp13, a viral helicase almost identical on SARS-CoV and SARS-CoV-2, was reported to bind SARS-CoV-2 G4s ^19^. Nsp3, another coronavirus protein, was shown to bind human host cell RNA G4s through the SARS-Unique Domain (SUD) of Nsp3 ^46^. Further work showed CNBP - a human cell protein that is elevated on infection and binds to SARS-CoV-2 genome - binds and unfold G4s from the SARS-CoV-2 genome *in vitro* ^5,20,47,48^. In addition, RG-1 region in the N gene of the SARS-CoV-2 was found to form a G4 structure that could be targeted using a G4 binding ligand ^30^. On the other hand, stabilization of a G4 within the mRNA of human TMPRSS2, which was elevated in the lungs of COVID-19 patients, inhibited TMPRSS2 translation leading to prevention of SARS-CoV-2 entry ^49^. Together these suggest the significance of RNA G4s, both viral and host, in the replication, transcription, and assembly of SARS-CoV-2 ^5,50^

CPZ and PCZ was implicated to be able to abrogate host cell infection by hepatitis C, dengue and some coronaviruses by inhibiting clathrin-mediated endocytosis ^51–55^. It must be also noted that CPZ and PCZ are used as antipsychotic and antidepressant drugs based to their effect on dopamine D2 and alpha-2 adrenergic receptors ^33^. A recent study further suggested antipsychotic drugs might be inhibiting viral replication ^56^. A clinical trial found that second-generation antipsychotics was associated with decreased risk of COVID-19 infection in patients in the New York State–wide psychiatric hospital system ^57^. Interestingly, though the decrease in risk of infection with CPZ (a first-generation antipsychotic) was not quite statistically significant, none of the 47 COVID-19 infected patients taking CPZ died of COVID-19 related causes. It is therefore likely that the effect of CPZ/PCZ against SARS-CoV-2 infection is through multiple mechanisms that affect both the host and the virus. The prophylactic and therapeutic activity of CPZ/PCZ observed by us against COVID-19 infection in hamsters (Figures 5 and 6) support this.

In conclusion, this is the first report showing FDA approved drugs CPZ and PCZ as potentially useful treatment options for COVID-19 due to its wide availability and relative affordability. We focused on targeting RNA structure in the form of G4s in the viral genome. Results demonstrate that secondary nucleic acid structures, that are relatively invariant (with low mutation frequency) in pathogenic genomes, could be effective targets against fast mutating strains like SARS-CoV-2.

## Materials and Methods

### Materials

Chlorpromazine and Prochlorperazine were purchased in the form of hydrochloride and dimaleate salt respectively, from Sigma (USA). All the RNA oligos were purchased from Sigma of HPLC-purified grade. The sequences of the oligonucleotides used in these studies are presented in Table-1. Concentration of oligonucleotides solution were determined from absorbance at 260 nm using molar extinction coefficient at 260 nm of 384.2 and 401.5 M^-1^ cm^-1^ for ORF1 WT and ORF1 Mut sequences respectively. Nuclease free water (DEPC untreated) was used throughout all the experiments. All biophysical experiments were done in 10 mM sodium cacodylate buffer (pH 7.4) containing 100 mM KCl at 25°C unless mentioned otherwise. The RNA oligos were heated at 95°C for 5 minutes and gradually cooled down to 25°C at the rate of 0.2 °C min^-1^ to facilitate quadruplex formation for the biophysical studies.

### Estimation of genome wide mutation frequencies and conservation analysis of pG4s

Genome wide mutation counts of globally circulating SARS CoV-2 genomes were retrieved from China National Center for Bioinformation (CNCB) (National Genomics Data Center, n.d.). This site provides the comprehensive details of over 29000 variation sites observed from about 5.4 million SARS CoV-2 genomes worldwide. The frequency of mutations at each nucleotide site were systematically estimated using Microsoft excel functions. In addition, the mutation frequency of the pG4 motifs was analyzed by estimating the average mutation frequency of each of the Guanines involved in G4 tetrad formation.

### Circular dichroism (CD) Spectroscopy

CD spectra experiments were carried out on JASCO J-815 spectropolarimeter equipped with a temperature-controlled cell holder and a cuvette with a path length of 1 cm. The oligos were used after going through the quadruplex formation procedure mentioned in the Materials section. Differential absorption spectra were recorded in the 200-300 nm range at room temperature. The represented spectrum is an automated average of three consecutive scans for each sample. CD titrations were carried out by the stepwise addition of CPZ or PCZ (up to 5 µM) to a cell containing 5 µM RNA oligo.

### Fluorescence assays

The assays were carried out on Horiba Scientific Fluoromax-4 spectrofluorometer at 25°C. For NMM treated samples, emission spectra were measured by using an excitation wavelength of 399 nm. The concentration of NMM and the RNA oligos was fixed at 1 μM. The oligos were used after going through the quadruplex formation procedure mentioned in the Materials section. For the displacement assay, the concentration of NMM and the RNA oligos was fixed at 1 μM while CPZ and PCZ were added in increasing molar equivalents.

### Isothermal Titration Calorimetry (ITC)

ITC measurements were carried out in a PEAQ-ITC titration calorimeter (Malvern Panalytical). Before loading, the solutions were thoroughly degassed. The oligos were used after going through the quadruplex formation procedure mentioned in the Materials section. The RNA oligo (15 μM) was kept in the sample cell, and 80 µl of CPZ/PCZ (1000 μM) dissolved in the same buffer (10 mM sodium cacodylate with 100 mM KCl) was filled in the syringe. Ligand solution was added sequentially in 3 µl aliquots (for a total of 13 injections, 6 s duration each) at 180 s intervals at 25°C. For CPZ, the integrated heat data were fit with one binding site model using the Microcal PEAQ ITC analysis software. For PCZ, due to the non-sigmoidal shape of the thermogram and the obvious presence of at least two independent binding processes, the thermogram obtained in ITC experiments was fit with two independent sites model in Microcal PEAQ ITC analysis software. Finally, the data was plotted using Origin pro 8.5 software.

### Reverse Transcription Inhibition Assay

Inspired from reverse transcriptase stalling used to identify RNA G4s, a reverse transcription assay from synthetic SARS CoV2 genomic RNA was done in presence of CPZ/PCZ (Kwok et al., 2016). Twist control 2, a synthetic SARS-CoV2 RNA template obtained from Twist Bioscience was used as the template RNA for this assay. The synthetic RNA template was reverse transcribed using Superscript II reverse transcriptase obtained from Invitrogen, in presence of 50μM of KCl. Reverse transcription was done using reverse primers such that ORF1 pG4-1 or a non G4 forming control region gets reverse transcribed (primer sequences below). Different concentrations of CPZ and PCZ were added in the reverse transcription reactions and compared against no drug treatment. The efficiency of the reverse transcription reactions was measured by quantifying the generated cDNA by qPCR using primers overlapping the ORF1 pG4 or the non G4 forming control region (primer sequences below). The same experiment was also done using RNA isolated from nasopharyngeal swabs of COVID-19 infected individuals following approvals by the Institutional Human Ethics Committee (IGIB)

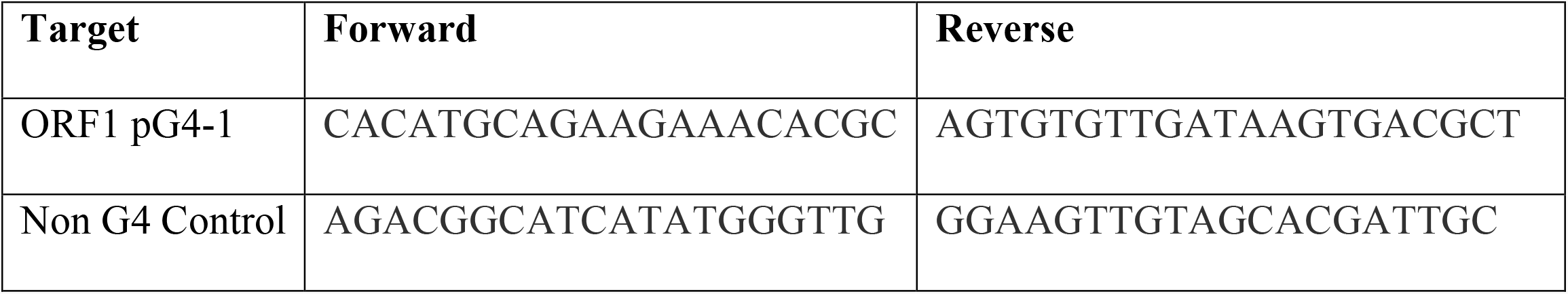

### Cell culture

All inhibition experiments were carried out on African green monkey kidney cells (Vero CCL-81). The cells were cultured in Dulbecco’s modified Eagle’s medium (DMEM, Gibco) supplemented with 10% heat inactivated Fetal Bovine Serum (Gibco) and 1 × Pen Strep (Gibco).

### Virus Generation

An isolate of B.6 strain of SARS-CoV-2 was used in this study. The isolation and characterization were described previously ^58^. In brief, viral transport media was first filtered through a 0.22 μm filter and was used as inoculum on Vero cells in a 96-well format. Upon the appearance of visible cytopathic effects (CPE), the viral culture supernatant was used to infect fresh cells to amplify the viral titer. The process was repeated until the supernatant displayed infectious titer in the range of 10^−7^ml^-1^.

### Cell infection and Pharmacological (G4 ligands) treatment

Vero cells were seeded in a 12-well format. At 90% confluency, cells were either mock-infected or infected with 1 MOI of SARS-CoV-2 in serum-free medium for 3 h in the presence of CPZ or PCZ. Subsequently, the viral inoculum was replaced with growth medium containing 10% FBS medium and CPZ/PCZ, or the vehicle controls (same volume either water or DMSO). After 24 h post infection (hpi), the media was replaced with fresh growth medium with the respective compounds, and cultured for another 24 h. At 48 hpi, the cell culture supernatants were collected to measure infectious particle count through PFU and extracellular viral RNA using quantitative real time PCR.

### Virus Titration (PFU)

The virus was titrated using PFU (Plaque-Forming Unit) assay in Vero cells following protocol mentioned in ^58^. Briefly Vero cells were seeded in 12-well plate at 90% confluency. The viral culture supernatant obtained after CPZ/PCZ treatment (mentioned above) was log-diluted from 10-1 to 10-7 in serum-free media and was added to a 100% confluent monolayer of Vero cells. Three hours post-infection the infection inoculum was replaced with agar media (one part of 2% low melting point agarose (LMA) mixed with one part of 2 × DMEM with 5% FBS and 1% Pen-Strep). 6–7 days post-infection, cells were fixed with 4% formaldehyde in 1 × PBS and stained with 0.1% crystal violet. The dilution which had 5–20 plaques was used for calculating PFU/ml. Each dilution was assayed in duplicate and the values from the duplicate wSells were averaged before calculating the respective PFU.

### RNA isolation and Real-time qPCR

RNA from cell culture supernatant after CPZ/PCZ treatment (mentioned above) was isolated using viral RNA isolation kit (MACHEREY-NAGEL GmbH & Co. KG). Real-time quantitative RT-PCR was performed in Roche LightCycler 480 using commercial kits. LabGun™ COVID-19 RT-PCR Kit was used to measure the RNA levels of SARS-CoV-2 RdRp and E genes while Fosun COVID-19 RT-PCR Kit was used to measure the RNA levels of SARS-CoV-2 ORF1ab and N genes following manufacturers’ protocol. Fold changes between samples were calculated by ΔΔ Cp method using the internal control (IC) for normalization.

### Cytotoxicity assay

The cytotoxicity of the drugs used in the study was determined using MTT((3-[4,5-dimethylthylthiazol-2-yl]-2,5-diphenyl) assay. Following the above-mentioned infection and CPZ/PCZ treatment, MTT was added to the cells at 24 hpi at final concentration of 0.5mg/mL in fresh media, following 6 h of incubation at 37°C, media was aspirated and formazan crystals were dissolved in 100μl of 100% DMSO, plates were incubated at RT for 20 mins with constant shaking and readings at 570nm were taken. A cell-free medium control was included to account for background due to phenol red in the medium.

### Animals

6-8 weeks old male golden Syrian hamsters were procured from CDRI and transported to small animal facility (SAF), THSTI and quarantined for 7 days. During the pre-treatment regime the animals were housed at small animal facility (SAF) and then were transferred to the Animal biosafety level-3 (ABSL-3) institutional facility for SARS-CoV-2 challenge study. The animals were maintained under 12 h light and dark cycle and fed standard pellet diet and water ad libitum. All the experimental protocols involving dosing and animal challenge were approved by institutional IAEC, IBS and RCGM.

### Virus generation for animal experiments

SARS-Related Coronavirus 2, Isolate USA-WA1/2020 virus was grown and titrated in Vero E6 cell line cultured in Dulbecco’s Modified Eagle Medium (DMEM) complete media containing 4.5 g/L D-glucose, 100,000 U/L Penicillin-Streptomycin, 100 mg/L sodium pyruvate, 25mM HEPES and 2% FBS. The stocks of virus were plaque purified at THSTI IDRF facility inside ABSL3 following institutional biosafety guidelines.

### SARS-CoV2 infection in golden Syrian hamster and dosing

Golden Syrian hamsters were randomly allotted to different drug groups (n=4), challenge control (n=2), remdesivir control (n=2) and unchallenged control (n=2) were housed in separate cages. The remdesivir group received a subcutaneous injection of remdesivir at 15 mg/kg body weight 1 day before and 1 day post infection ^39^. The pre-treatment group viz pCPZ & pPCZ started receiving 8mg/kg and 5mg/kg (respectively) of the drug through intraperitoneal administration each day starting from 3 days prior to the challenge and continued till end point (day 4 post infection). The therapeutic groups viz tCPZ and tPCZ received the drug 8 mg/kg and 5 mg/kg (respectively) through intra-peritoneal injection from the day of challenge for each day till the end point ^41,42^. All the animals, except unchallenged control, were challenged with 10^5^ PFU of SARS-CoV2 administered intranasally using a catheter while under anesthesia by using ketamine (150mg/kg) and xylazine (10mg/kg) intraperitoneal injection inside ABSL3 facility ^37,38,40^. Unchallenged control group received mock PBS intranasally. All the experimental protocols involving the handling of virus culture and animal infection were approved by RCGM, institutional biosafety and IAEC animal ethics committee.

### Gross clinical parameters of SARS-CoV2 infection

All infected animals were euthanized on 4 days post infection at ABSL3. Changes in body weight, activity of the animals were observed on each day post challenge. Post sacrifice, lungs and spleen of the animals were excised and imaged for gross morphological changes ^40^. Right lower lobe of the lung was fixed in 10% neutral formalin solution and used for histological analysis. The complete left lobe of the lung was homogenized in 2ml Trizol solution for viral load estimation. Spleen was homogenized in 2ml of Trizol solution. The tissue samples in trizol were stored immediately at −80 °C till further use.

### Viral load and Splenocytes qPCR

RNA was isolated from the lung and spleen samples using Trizol-choloroform method. Thereafter, RNA was quantitated by NanoDrop and 1 µg of total RNA was then reverse-transcribed to cDNA using the iScript cDNA synthesis kit (Biorad; #1708891) (Roche). Diluted cDNAs (1:5) was used for qPCR by using KAPA SYBR® FAST qPCR Master Mix (5X) Universal Kit (KK4600) on Fast 7500 Dx real-time PCR system (Applied Biosystems) and the results were analyzed with SDS2.1 software. A CDC-approved commercial kit was used for the quantification of the SARS-CoV-2 N gene for viral load estimation. The relative expression of each gene was expressed as fold change and was calculated by subtracting the cycling threshold (Ct) value of hypoxantine-guanine phosphoribosyltransferase (HGPRT-endogenous control gene) from the Ct value of target gene (ΔCT). Following primers were used ^40^.

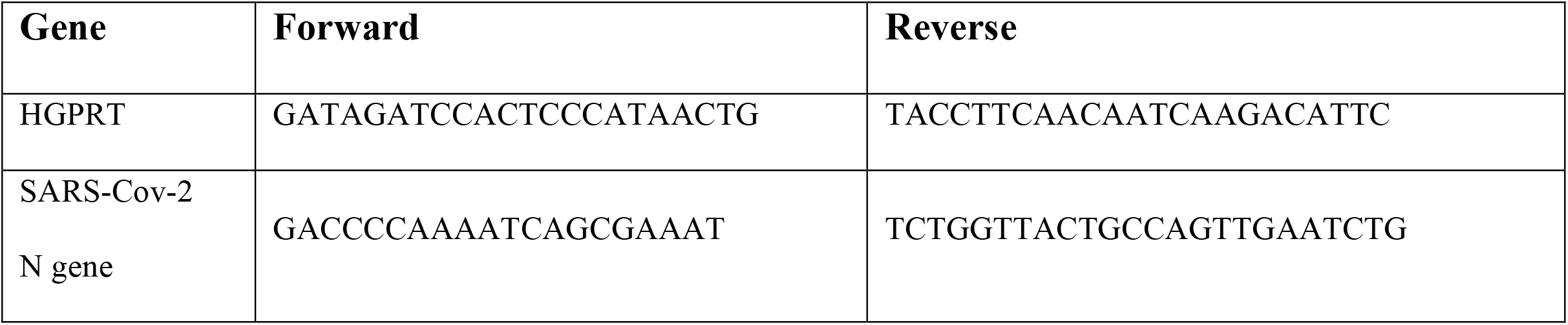

### Lung histology

Fixed lungs were processed and paraffin wax embedded blocks were transverse sectioned and stained with hematoxylin and eosin. The H & E-stained lung sections was then quantitatively examined under the microscope for pneumonitis, alveolar epithelial cell injury, inflammation, and lung injury on the scale of 0-5 by expert histologist. Images of the HE stained lungs sections were acquired at 40X magnification.

### Statistical analysis of data

p-values for the animal studies were calculated using ordinary one-way ANOVA for statistical significance. For all other experiments done in triplicates p-values were calculated using student’s *t*-test for statistical significance (*p < 0.05, **p < 0.01, ***p < 0.005).

## Acknowledgements

We are grateful to Sulochana Bagri, S.C. lab, for her help with proofreading the manuscript. This work was supported by research grants from Wellcome Trust/DBT India Alliance Fellowship (IA/S/18/2/504021) to S.C. Support from Council of Scientific and Industrial Research (CSIR) and Department of Biotechnology (DBT) to S.C. are also acknowledged.

## Author Contributions

S.C. conceived and directed the project. S.C. along with S.S.R. and S.Sharma designed the experiments. S.S.R. and M.R. analyzed the mutation frequency. S.S.R. and D.S. performed the biophysical experiments (CD, ITC, fluorescence turn-on/displacement). S.S.R. and P.S. performed the reverse transcription inhibition experiments. D.G. performed the Vero cell experiments. S.C. and Z.A.R. designed the hamster experiments. Z.A.R., S.Sadhu and M.R.T. performed the hamster experiments. S.Samal generated the virus for the hamster experiments. S.C., S.M., K.H.H., V.S., A.A., S.Sivasubbu and S.J. guided the experimental designs and manuscript writing. S.C., S.S.R. and S.Sharma wrote the manuscript. All authors read and commented on the manuscript.

## Competing interests

The authors declare no competing interests.

**Table S1:**
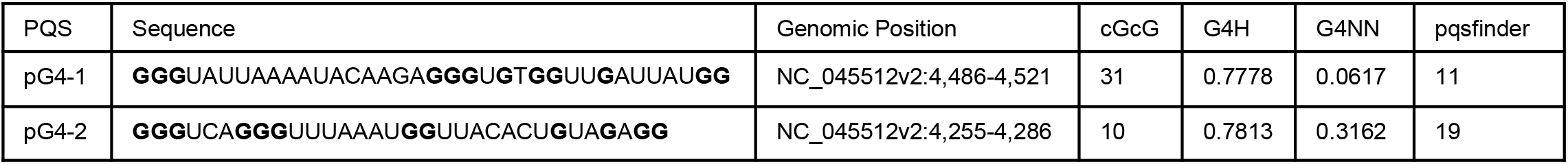
New pG4s identified in the SARS-CoV-2 genome that have bulges. Their sequences, genomic positions are mentioned along with the G4 propensity scores (cGcC-Consecutive G over consecutive C ratio, G4H-G4Hunter score, G4NN-G4 Neural Network, and pqsfinder score)

**Figure S1:**
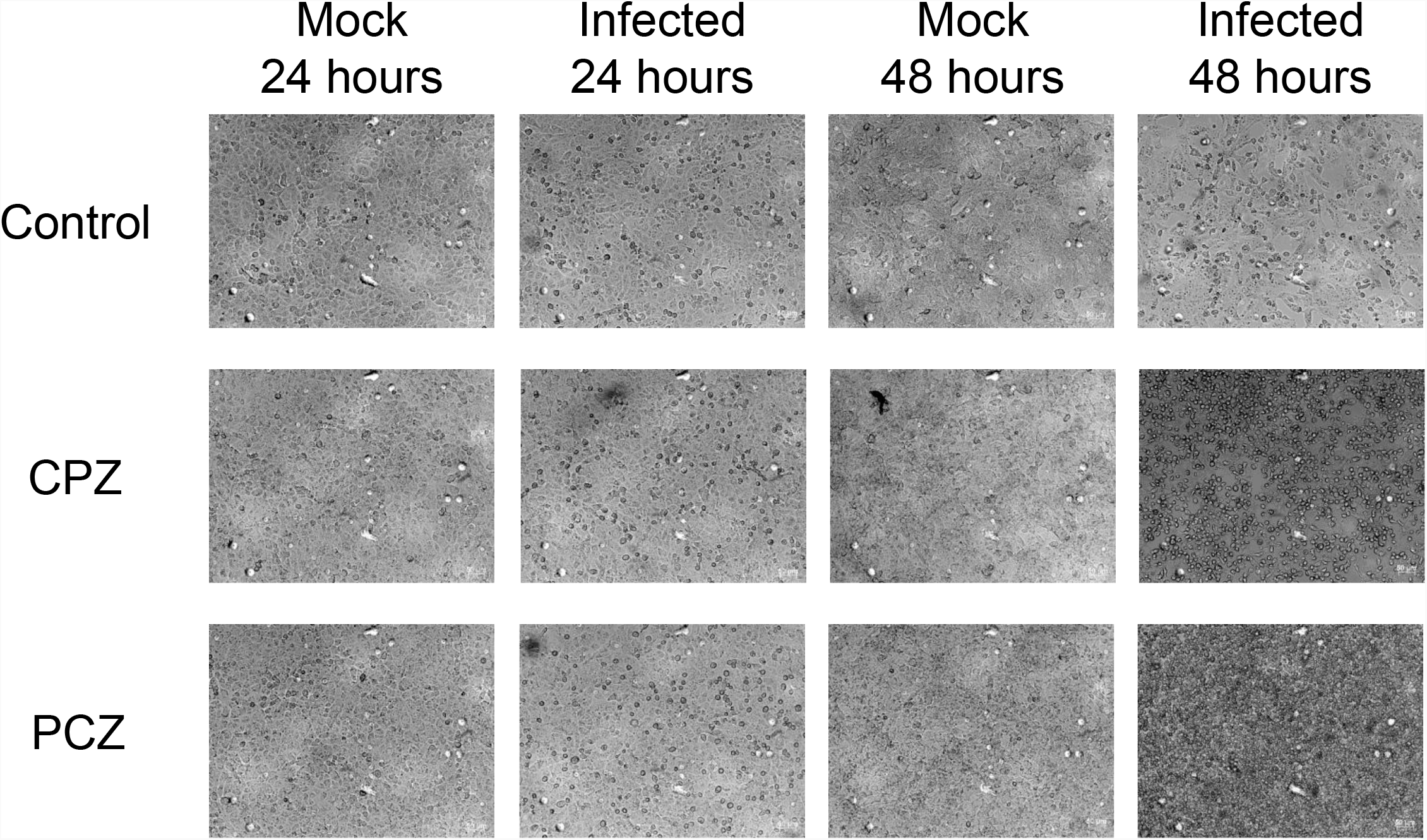
Brightfield images of Vero cells 24 or 48 hours after infection or mock treatment in the presence of CPZ, PCZ or no treatment control.

**Figure S2:**
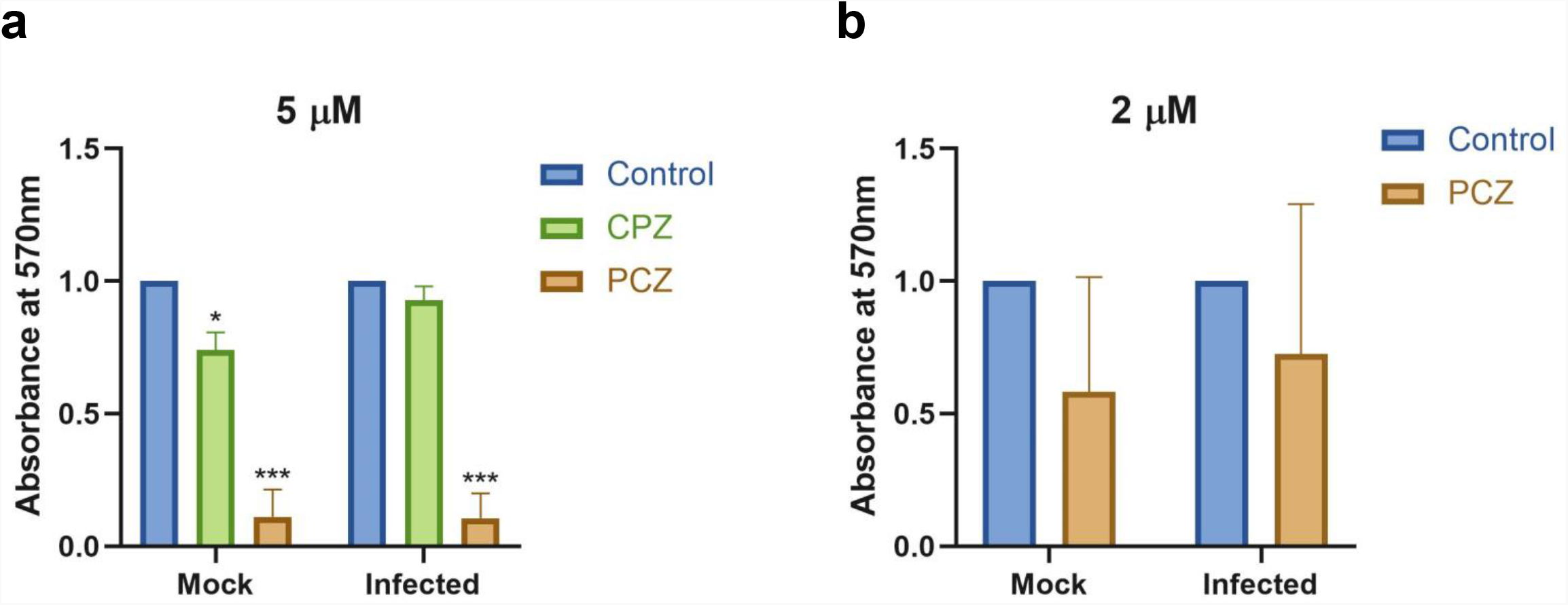
Viability of SARS-CoV-2 infected or uninfected Vero cells treated with 5 μM CPZ and PCZ (**a**) or 2 μM PCZ (**b**) compared against untreated control, measured by MTT assay. Mean ± SD (n=3); unpaired, two-tailed *t*-test.

**Figure S3:**
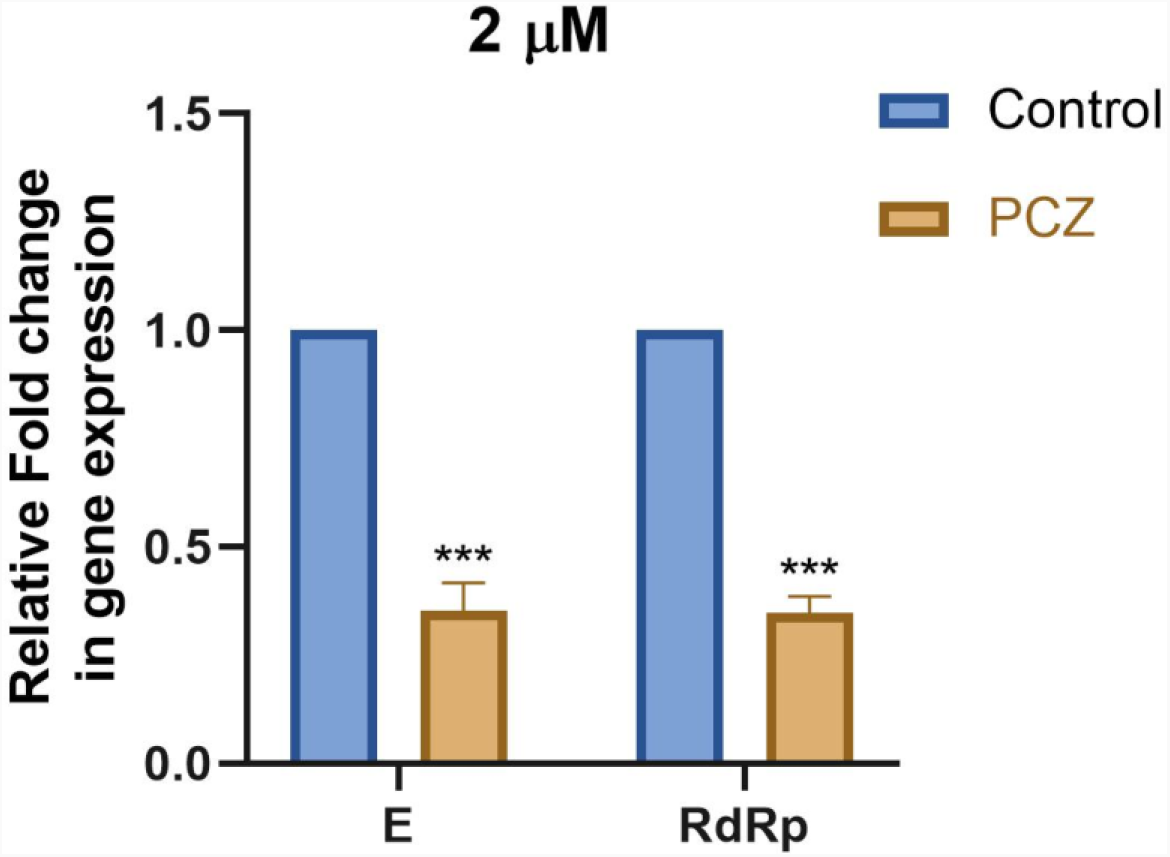
Expression of E and RdRp genes of SARS-CoV-2 in the extracellular media of infected Vero cells treated with 2 μM of PCZ compared against untreated control. Mean ± SD (n=3); unpaired, two-tailed *t*-test.

